# LiaX is a surrogate marker for cell-envelope stress and daptomycin non-susceptibility in *Enterococcus faecium*

**DOI:** 10.1101/2023.08.18.553907

**Authors:** Dierdre B. Axell-House, Shelby R. Simar, Diana Panesso, Sandra Rincon, William R. Miller, Ayesha Khan, Orville A. Pemberton, Lizbet Valdez, April H. Nguyen, Kara S. Hood, Kirsten Rydell, Andrea M. DeTranaltes, Mary N. Jones, Rachel Atterstrom, Jinnethe Reyes, Pranoti V Sahasrabhojane, Geehan Suleyman, Marcus Zervos, Samuel A. Shelburne, Kavindra V. Singh, Yousif Shamoo, Blake M. Hanson, Truc T. Tran, Cesar A. Arias

**Affiliations:** Division of Infectious Diseases, Department of Medicine, Houston Methodist Hospital, Houston, TX, USA; Center for Infectious Diseases, Houston Methodist Research Institute, Houston, TX, USA; Department of Medicine, Weill Cornell Medical College, New York, NY, USA; Center for Infectious Diseases, University of Texas Health Science Center, School of Public Health, Houston, TX, USA; Molecular Genetics and Antimicrobial Resistance Unit, Universidad El Bosque, Bogotá, Colombia; Department of Pathology, Microbiology and Immunology, Vanderbilt University Medical Center, Nashville, TN, USA; Department of Biosciences, Rice University, Houston, TX, USA; McGovern Medical School, University of Texas Health Science Center at Houston, Houston, TX, USA; Department of Infectious Diseases, Infection Control, and Employee Health, University of Texas MD Anderson Cancer Center, Houston, Texas, USA; Department of Internal Medicine, Division of Infectious Diseases, Henry Ford Hospital, Detroit, Michigan, USA; Department of Genomic Medicine, University of Texas MD Anderson Cancer Center, Houston, Texas, USA

**Keywords:** *Enterococcus faecium*, antibiotic resistance, daptomycin

## Abstract

Daptomycin (DAP) is often used as a first line therapy to treat vancomycin-resistant *Enterococcus faecium* (VR*Efm*) infections but emergence of DAP non-susceptibility threatens the effectiveness of this antibiotic. Moreover, current methods to determine DAP MICs have poor reproducibility and accuracy. In enterococci, DAP resistance is mediated by the LiaFSR cell membrane stress response system and deletion of *liaR* encoding the response regulator results in hypersusceptibility to DAP and antimicrobial peptides. The main genes regulated by LiaR are a cluster of three genes, designated *liaXYZ*. In *Enterococcus faecalis*, LiaX is surface exposed with a C-terminus that functions as a negative regulator of cell membrane remodeling and an N-terminal domain that is released to the extracellular medium where it binds DAP. Thus, in *E. faecalis*, LiaX functions as a sentinel molecule recognizing DAP and controlling the cell membrane response, but less is known about LiaX in *E. faecium*. Here, we found that *liaX* is essential in *E. faecium* (*Efm*) with an activated LiaFSR system. Unlike *E. faecalis*, *Efm* LiaX is not detected in the extracellular milieu and does not appear to alter phospholipid architecture. We further postulated that LiaX could be used as a surrogate marker for cell envelope activation and non-susceptibility to DAP. For this purpose, we developed and optimized a LiaX ELISA. We then assessed 86 clinical *E. faecium* BSI isolates for DAP MICs and used whole genome sequencing to assess for substitutions in LiaX. All DAP-R clinical strains of *E. faecium* exhibited elevated LiaX levels. Strikingly, 73% of DAP-S isolates by standard MIC determination had elevated LiaX ELISAs above the established cut-off. Phylogenetic analyses of predicted amino acid substitutions showed 12 different variants of LiaX without a specific association with DAP MIC or LiaX ELISA values. Our findings also suggest that many *Efm* isolates that test DAP susceptible by standard MIC determination are likely to have an activated cell stress response that may predispose to DAP failure. As LiaX appears to be essential for the cell envelope response to DAP, its detection could prove useful to improve the accuracy of susceptibility testing by anticipating therapeutic failure.

## INTRODUCTION

Vancomycin-resistant enterococci (VRE) are organisms that cause serious healthcare-associated infections (1). *Enterococcus faecium* (*Efm*) account for greater than 75% of VRE (2) and a concerning number develop resistance to additional antibiotics, exhibiting the multidrug-resistant phenotype. Daptomycin (DAP), a cyclic bactericidal lipopeptide, has become a first line choice to treat severe VRE infections. The mechanism of DAP action involves insertion in the membrane of the calcium-bound antibiotic molecule, where it interacts with phosphatidylglycerol and lipid II intermediates to disrupt cell wall biosynthesis and cell membrane homeostasis (3). However, after introduction of DAP in clinical practice, resistance (DAP-R) has been increasingly documented. Indeed, some data suggest that DAP-R occurs in up to 20-40% of VRE bloodstream infection (BSI) isolates in some institutions (4, 5) and can emerge during therapy (6, 7), jeopardizing the clinical utility of this antibiotic against recalcitrant VRE infections. Moreover, DAP-R has also been documented in isolates recovered from patients who have never been exposed to the antibiotic (8).

The LiaFSR system is a three-component cell envelope stress regulatory system that has been implicated in DAP resistance both in *Enterococcus faecalis* and *Efm*. Deletion of the genes encoding the response regulator LiaR in both species has been shown to cause hypersusceptibility to DAP (9, 10). Characterization of the LiaR regulon in both *E. faecalis* and *Efm* indicated that the main target of the response regulator is a cluster of genes encoding three proteins designated LiaX, LiaY and LiaZ (11, 12). LiaX was initially characterized as a protein that binds PBP-5 in *Efm* and is likely involved in β-lactam resistance (13). Subsequently, in *E. faecalis*, it was found that LiaX is a surface-exposed protein harboring two distinct domains with specific and independent functions. Indeed, the C-terminal domain of LiaX functions as the actual cell membrane effector of LiaFSR by negatively controlling phospholipid remodeling that occurs during the “attack” by DAP and antimicrobial peptides, a phenotype that is an indicator of cell membrane adaptation in *E. faecalis*. On the other hand, the N-terminus appears to be released from the surface during cell envelope stress and functions as a sentinel signal transduction molecule by binding to DAP and antimicrobial peptides, a complex that is likely to be recognized on the cell surface to maintain the cell membrane stress response, critical for cell survival (12). In *Efm* however, the gene organization of the *liaXYZ* cluster varies and, most importantly, the major membrane changes in anionic phospholipid microdomains associated with development of DAP resistance by *E. faecalis* do not seem to occur in *Efm*.

Establishing DAP susceptibility in clinical strains of *Efm* has been challenging, mainly because choosing a breakpoint is complicated by the wild-type distribution of the isolates with many exhibiting higher MICs than *E. faecalis* and falling beyond the breakpoint initially proposed. This situation has prompted CLSI to change the DAP breakpoints (14) and establish different values for *E. fa*ecalis and *Efm*. Moreover, DAP MIC determination is influenced by many variables including the calcium concentration and the specific lot of Mueller-Hinton used by clinical laboratories. Further, reproducibility of broth microdilution and available Etests is questionable, with data suggesting poor interlaboratory agreement when performing MIC of the same isolates (15). In this scenario, the use of DAP for severe VRE infections becomes difficult, particularly when susceptibility testing is not accurate and its reliability is dubious, putting at risk patients who are often immunocompromised and prone to poor outcomes. While CLSI refers to *Efm* strains with DAP MICs ≤ 4 μg/ml as susceptible dose-dependent (SDD), we will use the term susceptible for ease of presentation.

In this work, we hypothesized that LiaX levels could be used to better assess DAP susceptibility in clinical *Efm* strains compared to standard MIC determination. Since standard susceptibility testing for DAP has major limitations in reproducibility and consistency (15), detection of a surrogate marker of cell membrane adaptation that accurately reflects susceptibility to DAP in *Efm* could serve as a more reliable tool to detect DAP resistance or tolerance and may potentially be clinically useful. During this study, we also provide further insights into the role of *liaX* in the cell envelope stress response of *Efm*.

## RESULTS

### LiaX in E. faecium is not detected in the extracellular milieu

In *E. faecalis*, LiaX is readily identified in the extracellular milieu, which is thought to bind to incoming calcium-bound DAP molecules and antimicrobial peptides, as a mechanism for sensing incoming “threats”. Thus, we sought to investigate if LiaX could also be readily detected in the extracellular medium in *Efm*. To characterize the localization and dynamics of LiaX in *Efm*, we generated anti-LiaX antibodies for use in immunoblotting and enzyme-linked immunosorbent assays (ELISA). We overexpressed full-length LiaX in *E. coli* from the well-characterized DAP-R *Efm* reference strain R494 (16). Subsequent purification resulted in an aliquot of LiaX protein assessed to have a purity of >95% (Fig. S1). Using the purified LiaX protein, we raised anti-LiaX antibodies in goats, which subsequently underwent high-affinity purification against LiaX.

To assess the analytical specificity of the purified polyclonal antibodies (Ab), immunoblotting was performed on *Efm* strains which harbored deletions of *liaR* and/or *liaX.* We generated mutants of *liaR* (the gene encoding the response regulator of the LiaFSR system) in a commensal strain of *Efm* (TX1330RF) (17). Deletion of *liaR* (TX1330RFΔ*liaR*) is expected to abolish transcription of the *liaXYZ* gene cluster (9–12). Furthermore, we attempted to delete *liaX* in TX1330RF but were only able to delete this gene in the background of TX1330RFΔ*liaR*, generating a double mutant (TX1330RFΔ*liaRΔliaX*) (see below). Using these strains, we were able to detect bands at the expected molecular weight of *Efm* LiaX (58.6 kDa) with purified LiaX and cell lysates of TX1330RF and TX1330RFΔ*liaR*, but not with TX1330RFΔ*liaRΔliaX*, indicating robust specificity of the primary polyclonal antibody (Ab) for LiaX (Fig. 1A). These results also suggested that a basal amount of LiaX is produced even in absence of the LiaR response regulator.

**Figure 1.**
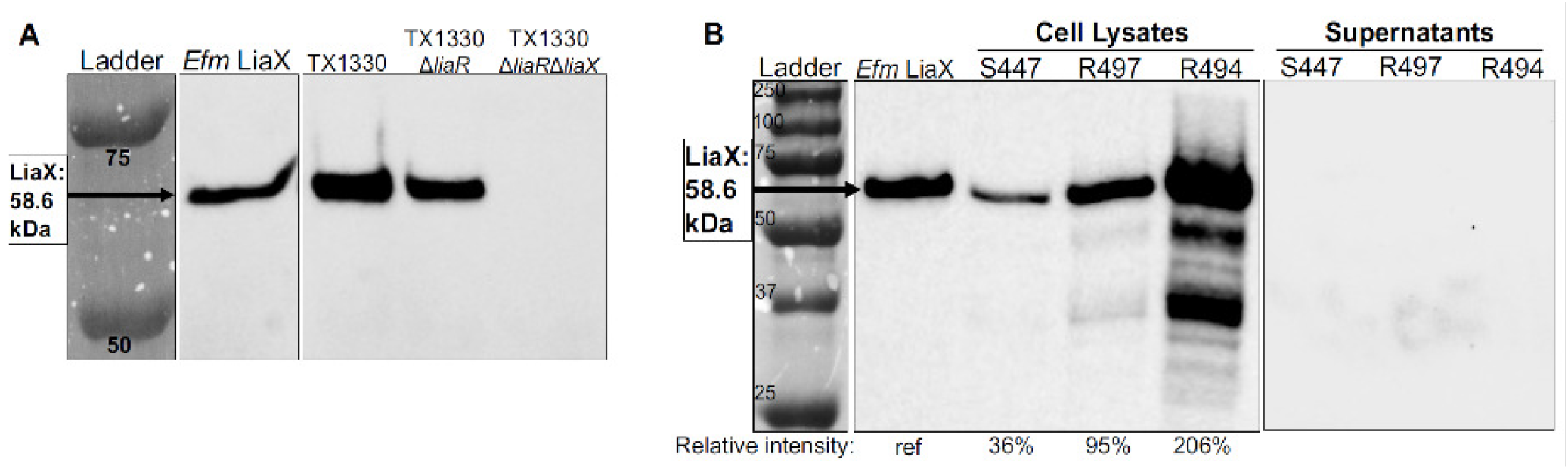
lmmunoblots of *Efm* Liax protein, mutant *Efm* strains, and clinically-derived *Efm* Isolates. **A)** We were able to detect Liax in TX1330Δ*liaR* even In the absence of LiaR. The Liax signal Is abolished In TX1330Δ*liaR*Δ*liax*. lmmunoblots of cell lysates of TX1330 and derivative strains show bands present at the expected molecular weight of *Efm* LiaX (58.6 kDa). **B)** LiaX Is detected In higher amounts in DAP-R compared to OAP-S clinically-derived *Efm* isolates and Is not detected in extracellular media. lmmunoblots of cell lysates and protein-precipitated extracellular media (supernatants) demonstrate Increased LiaX production In DAP-R R497 and R494 compared to DAP-S S447 cell lysates. Supernatants of the same strains showed not detectable LiaX. 2 ng of purified *E. faecium* LiaX protein was used as a positive control. The numbered ladder Indicates the molecular weights of bands in kilodaltons (kDa). Relative band Intensities are shown below the clinically-derived strains for the 58.6 kDa band In cell lysates. Band Intensities were normalized to positive control purified *E. faecfum* LiaX.

Next, using the specific anti-LiaX antibodies, we sought to investigate the localization of LiaX in *Efm* isolates. Of note, in clinical and laboratory strains of *E. faecalis*, LiaX can be detected on the cell surface and in the extracellular media (N-terminal domain). Immunoblotting was performed on well-characterized clinical strains of DAP-S (S447) (6, 16, 18) and DAP-R *E. faecium* (R497 and R494) (16) using purified polyclonal goat antibodies. Strains were grown to mid-exponential phase, then cells were separated from media for further analysis. Cells were lysed by glass beads, while the media underwent filter sterilization, and proteins were precipitated by the addition of trichloroacetic acid (TCA). Immunoblotting was performed on all samples after normalization for protein concentration. Quantification of the LiaX protein in cell lysates from DAP-R R497 and R494 was performed using Western blot analysis. Cell lysates from R497 and R494 exhibited higher intensities for the bands corresponding to LiaX, with values of 95% and 206%, respectively, relative to the intensity of the positive control. In contrast, the cell lysate of DAP-S strain S447 showed an intensity only 36% of the positive control. Most importantly, we were unable to detect LiaX in the supernatants of any strain, suggesting that, unlike *E. faecalis*, LiaX in *E. faecium* is not released to the extracellular milieu (Fig. 1B).

### Performance and optimization of the LiaX ELISA in clinical strains of E. faecium

We then optimized sensitivity of the purified polyclonal Ab for use in an ELISA assay by conducting a dilution titer. The purified Ab resulted in higher *A*_405nm_ at every Ab dilution compared to unpurified Ab. Simultaneously, there was absence of off-target or “background” signal (*A*_405nm_= 0) from the goat pre-immune sera (Fig. 2), indicating the absence of immunoreactivity to LiaX prior to immunization with LiaX. The ELISA titer was performed with a dilution of commercial enzyme-conjugated secondary Ab recommended by the manufacturer, which had previously been optimized in enterococcal bacterial studies (1:2000 or 0.3 μg/mL) (19). The titer was also performed with a constant antigen concentration of 10 ng of LiaX protein, which was representative of the *A*_405nm_ that was observed from ELISA performed on *Efm* strains in preliminary studies. The purified primary Ab dilution to produce the highest and most discriminatory signal for ELISA was determined to be 1:400 ([Ab] = 4.185 μg/mL).

**Figure 2.**
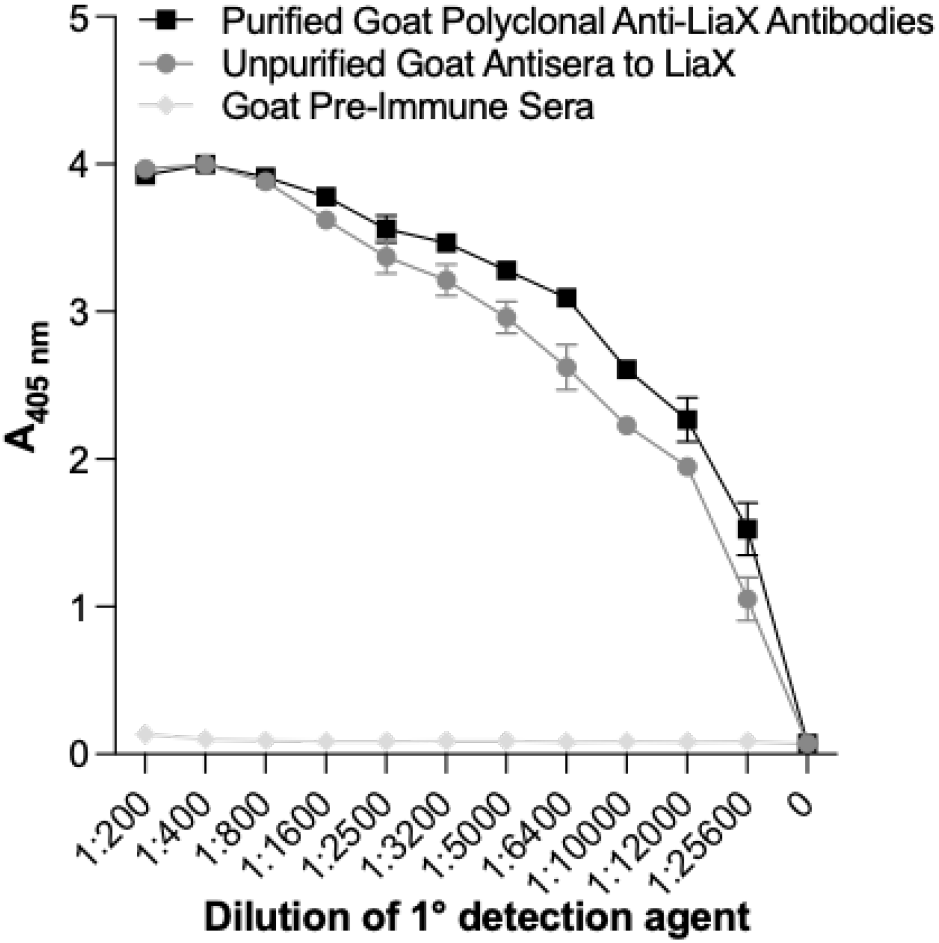
Titer of multiple primary detection agents against LiaX. ELISA assay with 10 ng *Efm* LiaX antigen coated per well. Results are shown in triplicate, plotted with mean and SEM. Not all error bars are visible due to small size.

We subsequently did a checkerboard titration of primary Ab and enzyme-conjugated secondary Ab using four different *Efm* strains (DAP-S strains S447 and 503, and DAP-R strains R446 and 1547). These strains were known to yield *A*_405nm_ values for the LiaX ELISA from the minimum to maximum spectrophotometer detection limits of visible assay absorbance from preliminary studies (Fig. S2). The dilutions that were found to yield the *A*_405nm_ values within the optimal accuracy range of our spectrophotometer with the greatest resolution were 1:400 for primary Ab and 1:2000 for secondary Ab. Using these concentrations, there was no off-target detection in the blank controls, indicating a highly specific assay.

After optimization by systematic checkerboard titration, the ELISA was performed on a collection of previously well-characterized, whole-genome-sequenced, clinically-derived *Efm* (20) with a wide range of DAP MICs (Table S1). The LiaX ELISA was able to differentiate between DAP-R and DAP-S strains using a preliminary cutoff at the *A*_405nm_ level based on DAP-S strain S447 (Fig. 3). Assays of reference strains were performed in four different iterations (4 different microtiter plates) over a 6-month period, with intra-assay coefficient of variation (CoV) ranging from 0.2% to 9.0%, and overall inter-assay CoV ranging from 7.4% to 23.8% (Table S2). These observations fell within our pre-determined range of acceptable variation (highest CoV of 20-25%) for the LiaX ELISA.

**Figure 3.**
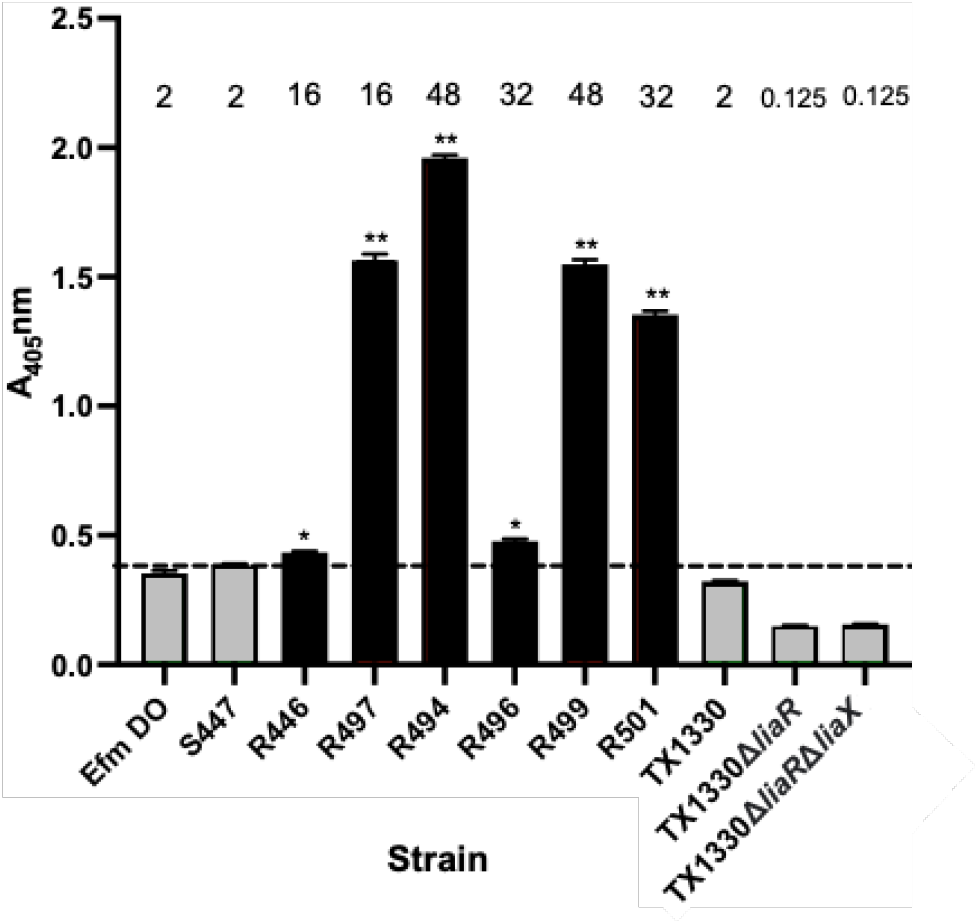
Whole-cell indirect LiaX ELISA of *Efm* reference and deletion strains using goat purified polyclonal Ab. All DAP-R strains (black boxes) have higher LiaX levels than DAP-S strains (gray boxes). *E. faecium* TX1330 with a double deleltion of *liaX* and *liaR* does not show any difference in LiaX detection compared to TX1330, which has only a deletion of *liaR.* DAP MICs are shown at the top of the chart for each strain. The dotted line indicates the maximum value of the highest A_405nm_ the DAP-S strain S447, *p<0.05, .. p<0.001 compared to S447.

Once the validation step was carried out in well-characterized *Efm* strains, we performed the ELISA LiaX test on a collection of 86 clinical strains of *Efm*. The *Efm* isolates had been collected from a multicenter global cohort of patients with enterococcal bacteremia – the Vancomycin-Resistant Enterococcal Bloodstream Infection Outcomes Study (VENOUS) (21). We chose the isolates based on the availability of whole genome information. We performed standardized gradient strip and broth microdilution assays (BMD) to determine DAP MICs on all 86 *Efm* clinical isolates. DAP MICs were overall higher when performed by gradient strip compared to BMD (by a median 2-fold higher, IQR 1.5-3). Of note, there was no categorical disagreement in MICs between gradient strip and the BMD method using the current CLSI interpretations (data not shown). DAP MICs determined by BMD were plotted against the LiaX ELISA *A*_405nm_ on a receiver operating characteristic (ROC) curve, and the cutoff was determined by maximizing the Youden index (22). Of note, a high proportion of *Efm* isolates (n=64; 74%) in the VENOUS cohort exhibited an elevated LiaX ELISA *A*_405nm_ using the established cutoff (Fig. 4). We only had 3 *Efm* isolates with confirmed MICs within the resistance range (≥ 8 µg/mL) and all exhibited a LiaX ELISA well above the cut-off value. Most importantly, we had a total of 61 out of 83 (73.5%) *Efm* isolates with DAP-S MICs (DAP MIC ≤ 4 µg/mL) that exhibited LiaX ELISA *A*_405nm_ values above the cut-off. When categorizing the results of the ELISA by MIC determined by BMD, more isolates with MIC < 2 µg/mL had the LiaX ELISA *A*_405nm_ above the cut-off compared to isolates with MICs ≥ 2 – 4 µg/mL (Fig. 4). This situation was reversed when using gradient strips with more isolates with MICs ≥ 2 – 4 µg/mL exhibited a high LiaX ELISA *A*_405nm_ compared to those with MICs < 2 µg/mL.

**Figure 4.**
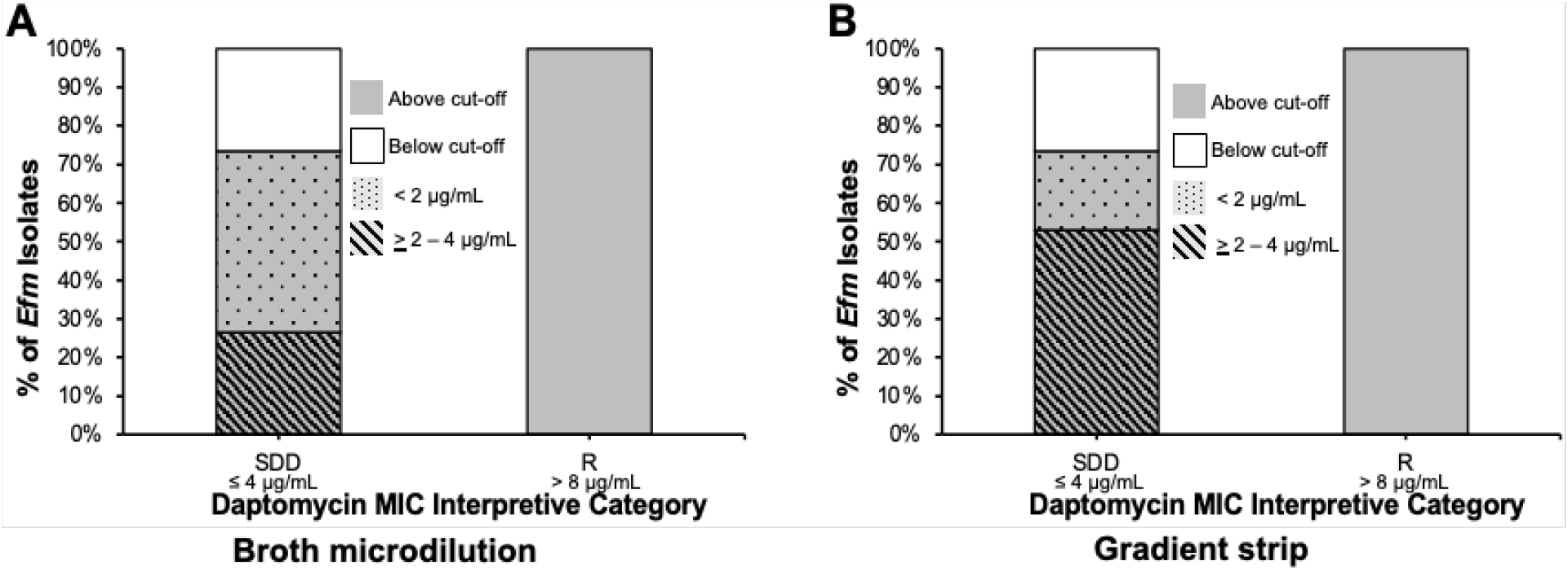
Detection of LiaX by ELISA in daptomycin-resistant and susceptible *Efm.* Minimum inhibitory concentrations (MICs) by botih **A)** broth microdilution and **B)** gradient strip tests.

Next, we evaluated the LiaX amino acid sequences of the 86 clinical isolates and 8 clinical strains used as references to assess sequence variations that may affect the performance of the LiaX ELISA. The LiaX amino acid sequences were compared to the consensus sequence of *Efm* TX16 (“*Efm* DO”). The majority (59%) of the *Efm* clinical isolates had a LiaX amino acid sequence with 100% identity to that of *Efm* DO. There were 11 other variants of LiaX, with changes encompassing 1 to 11 amino acid substitutions (Table S3) compared to the consensus sequence. There were no non-sense or frameshift alterations detected. Phylogenetic analysis of the LiaX variants showed a main cluster of LiaX variant 1 through 7, and a separate cluster of LiaX variant 8 through 12 (Fig. 5). We did not find any statistically significant association between any particular LiaX variant and discrepant DAP MIC/LiaX ELISA (i.e. DAP-S MIC with elevated LiaX ELISA) results. These findings suggest that the LiaX amino acid sequences identified in this study do not affect binding of the primary anti-LiaX antibody in the LiaX ELISA and, therefore, performance of the assay. Additionally, no association was found between LiaX variants and DAP MICs or LiaX ELISA *A*_405nm_ overall.

**Figure 5.**
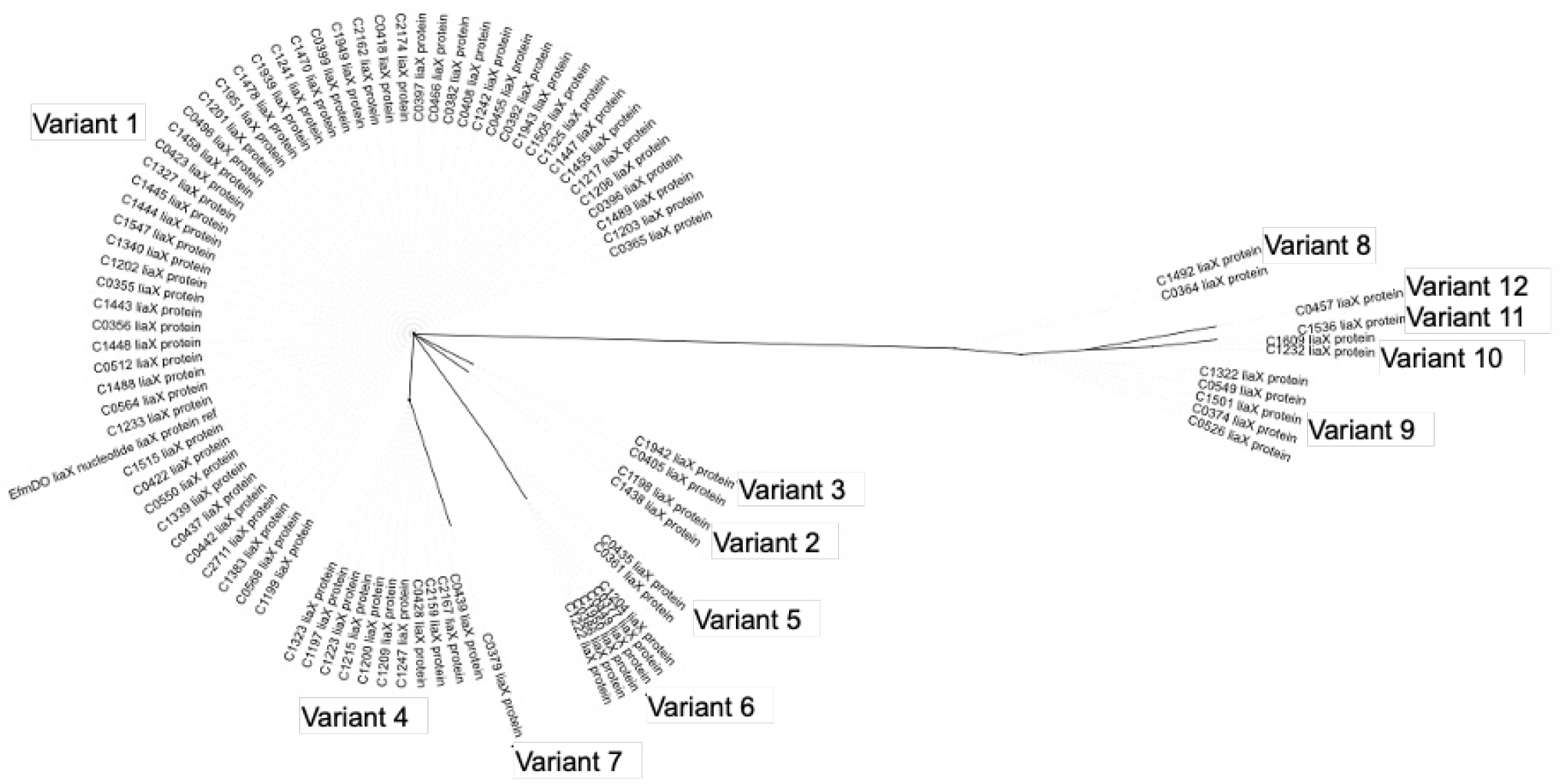
Phylogenetic analysis of LiaX sequences from *E. faecium* using a maximum likelihood tree.

### LiaX is essential in *E. faecium* in strains with an intact LiaFSR syste*m*

In *Efm*, LiaX is a 522-amino acid protein similar in primary sequence to LiaX of *E. faecalis*, harboring an N-terminal domain predicted to be mainly α-helices and a C-terminus composed of β-pleated sheets (Fig. S3). To gain insights into the LiaR regulon in *Efm*, we had previously (11) performed a transcriptional mapping of genes regulated by LiaR in the *Efm* DAP-R clinical strain R497 and its *liaR* deletion mutant derivative (R497Δ*liaR*). As mentioned above, we expected that deletion of *liaR* would abolish *liaX* expression. Indeed, the genes of the *liaXYZ* operon were downregulated in the R497Δ*liaR* mutant (11). We confirmed these results by qRT-PCR, which demonstrated downregulated expression of *liaFSR/liaXYZ* in the *liaR* deletion mutant (Fig. S4), supporting the notion that *liaXYZ* were under the control of LiaR in *Efm* clinical strains.

Subsequently, we attempted to delete *liaX* in the clinical strain DAP-R R497. Several attempts for mutagenesis were unsuccessful. We then used a different strain of *Efm* (TX1330RF), a clade B isolate that is a gut colonizer from the gastrointestinal tract of a human (17), which we have previously used for targeted mutagenesis (23, 24). Of note, TX1330RF is a DAP-susceptible, rifampin and fusidic acid-resistant derivative of TX1330. Using a markerless system (25), we attempted deletion of *liaX*, and although we obtained first-event recombination integrants, we were unable to obtain a null *liaX* mutant. After multiple attempts using different strains employing both the PheS* counterselection and CRISPR/Cas9 systems, we were unable to recover any *liaX* mutants in TX1330RF.

As manipulation of *liaX* proved challenging in both DAP-S and DAP-R *E. faecium* strains, we hypothesized that LiaX was an essential protein and that a basal level of this protein was necessary to maintain cell envelope integrity and response to cell surface stressors through the action of the LiaFSR system. Thus, we postulated that deletion of *liaX* might be possible in strains which have a non-functional LiaFSR system. This notion was supported by the fact that we had previously deleted *liaR* in several *Efm* strains (9, 10). Thus, we generated a *liaR* deletion mutant in TX1330RF prior to attempting *liaX* mutagenesis. Using this strategy, we were able to create a *liaX* deletion mutant in the TX1330RFΔ*liaR* background (TX1330RFΔ*liaR*Δ*liaX*). Since TX1330RFΔ*liaR* is DAP-hypersusceptible (MIC=0.125) compared to its parent TX1330RF (Table S1), a pattern previously observed in other *Efm* strains upon deletion of *liaR* (9), deletion of *liaX* did not have any effect on the DAP MIC (Table S1), supporting the assumption that LiaX is only essential within the context of a functional LiaFSR system.

Next, we attempted to complement *liaR* into TX1330RFΔ*liaR*Δ*liaX* to obtain a strain which functionally had only a *liaX* deletion. Efforts to complement *liaR* in TX1330RFΔ*liaR*Δ*liaX* in its native chromosomal location were unsuccessful, indicating that an active *liaR* affects cell viability in the absence of LiaX. Ultimately, we attempted a conditional complementation of *liaR* using the nisin-controlled vector pMSP3535 and successfully generated a derivative strain [TX1330RFΔ*liaR*Δ*liaX* (pMSP3535::*liaR*)] that was viable. However, the strain showed a major delay in growth of ∼3 hours compared to its parental TX1330RF (Fig. S5), supporting the notion that the absence of *liaX*, in the background of a functional LiaFSR system, causes major biological fitness costs. These results also suggest that LiaX in *Efm* is important for cell viability and that it functions as a major modulator of the LiaFSR response.

### Cell membrane phospholipid architecture and LiaX ELISA of E. faecium liaX and liaR mutants

In *E. faecalis*, activation of the cell envelope response to DAP and antimicrobial peptides via the LiaFSR system is accompanied by major changes in anionic phospholipid distribution (12, 26). Specifically, anionic phospholipid microdomains migrate away from the septum in an attempt to divert positively charged DAP from critical septal areas and prevent major damage during cell division and viability. Using NAO staining, we have previously (9) evaluated the architecture of anionic phospholipid microdomains in clinical strain pairs of DAP-S and DAP-R *Efm* and found that development of DAP-R is not associated with changes in phospholipid redistribution. As expected, deletion of *liaR* or both *liaX* and *liaR* in TX1330 also did not produce any changes in phospholipid microdomain distribution compared to parental wild-type strain (Fig. S6).

Finally, using our LiaX ELISA assay, we evaluated LiaX presence in the above mutants. Deletion of *liaR* in *Efm* strain TX1330RF (TX1330RFΔ*liaR*) led to a decrease in the *A*_405nm_ of the LiaX ELISA to a basal level compared to the parental (Fig. 3). Deletion of *liaX* in TX1330RFΔ*liaR* did not lead to any further decrease in the *A*_405nm_ of the LiaX ELISA, suggesting that the ELISA signal is likely due to background “noise”.

## DISCUSSION

DAP is considered a bactericidal first line antibiotic against vancomycin-resistant *Efm* despite lacking FDA indication for this purpose, particularly in highly immunosuppressed patients where bactericidal therapy is often preferred. A major issue of using DAP against vancomycin-resistant *Efm* is the emergence of resistance during therapy. Furthermore, there is evidence that DAP resistance can emerge even in the absence of DAP exposure (8), suggesting that the mechanism of protection against the antibiotic likely involves a major strategy that responds to DAP and other cell envelope stressors, including antimicrobial peptides. Indeed, the LiaFSR system appears to be a “broad” cell envelope stress system that can be readily activated in the presence of antibiotics. The LiaFSR system of *Efm* has similar target genes as those of *E. faecalis*, namely *liaXYZ*. However, there are major differences in the mechanistic strategy underlying DAP-R between *E. faecalis* and *Efm*. Indeed, the main global strategy to counteract the effect of DAP in *E. faecalis* appears to be to divert the antibiotic from the septum to other areas of the cell membrane. This phenomenon involves major changes in the cell envelope, including redistribution of anionic phospholipid microdomains (26). In contrast, *Efm* seems to “repel” DAP from the cell surface in a mechanism that likely involves changes in surface charge without major alterations in phospholipid architecture (20, 27). Regardless of the mechanism, the LiaFSR system-mediated response seems to be a key factor to successfully survive in the presence of DAP in both enterococcal species.

LiaX emerges as a critical effector of the LiaFSR system playing a key role at the cell membrane and envelope level. This protein was initially described (13) in *Efm* bound to penicillin-binding protein 5 (PBP5) (designated Pbp5-associated protein; P5AP). Indeed, P5AP (LiaX) was found to be a mediator of cephalosporin resistance in *Efm* whose typical class A PBPs are not functional. Moreover, the expression of *p5ap* (*liaX*) was noted to be influenced by the activity of the serine-threonine phosphatase/kinase system (13). Consistent with earlier studies and our transcriptional and ELISA experiments, the main regulator of *liaX* is LiaR. However, other factors could influence its expression since we were able to detect LiaX even in absence of a functional LiaR regulator, confirming those initial observations that this key protein can be regulated by different molecular systems. Although the function of LiaX in *Efm* is not fully understood, its association with PBPs and our mutagenesis experiments suggest that *liaX* is essential during cell envelope stress and indicates an even broader role for LiaX in adapting the cell membrane to DAP.

We have previously shown (12) that, in *E. faecalis*, DAP-R clinical isolates exhibited elevated LiaX levels, as detected by ELISA. Moreover, determination of DAP MICs using standard methodologies seem to be inaccurate and not reproducible between clinical microbiology laboratories, even in settings with known expertise in performing MIC determinations (15). Furthermore, there are well documented cases of failure of DAP monotherapy (7, 28) in patients whose isolates were reported “susceptible”. The challenges in DAP MIC testing are derived from the fact that variations are observed in the media lots and calcium concentrations, making the standardization of the test very difficult (29). Therefore, novel tools that could predict susceptibility to DAP more accurately are clearly needed.

Based on our initial observations and under the hypothesis that LiaX is a key protein in the presence of cell-envelope targeting antimicrobials in enterococci, we evaluated the possibility that LiaX may be used as a more accurate surrogate marker of cell envelope stress and, therefore, DAP susceptibility in *Efm*. Thus, we first developed and optimized an ELISA method for LiaX, assessing its performance on well-characterized strains of *Efm*. Consistent with our hypothesis, DAP-R strains indeed had significant higher levels of LiaX compared to DAP-S clinical strains (similar to *E. faecalis*). Further, using an established cut-off that resulted from initial experiments, we could identify all DAP-R strains with MIC ≥ 8 µg/ml. Subsequently, to validate our findings, we applied our LiaX ELISA method to 86 *Efm* isolated from bacteremia recovered from patients in an ongoing large cohort study of enterococcal bacteremia (VENOUS) (21). The isolates were selected based on a convenience sample from sites in Houston and Detroit participating in VENOUS and mostly recovered from patients with hematological malignancies and solid organ transplant recipients.

As expected, all DAP-R isolates within our validation cohort with MICs ≥ 8 µg/mL exhibited elevated LiaX levels supporting the correlation between an upregulated cell membrane stress and DAP non-susceptibility. Most importantly, 73% of *Efm* reported as “susceptible dose-dependent” exhibited LiaX levels above the established cut-off. This finding suggests that a large majority of invasive *Efm* (all VENOUS isolates were recovered from the blood of included patients) have already triggered membrane adaptation changes that are likely to affect DAP susceptibility. The “stressors” are likely to include antimicrobial peptides produced by the innate immune system during the process of gut colonization and infection within the human host. These findings also provide some explanation of why some *Efm* strains that had never been exposed to DAP are non-susceptible at the first encounter with the antibiotic.

The discrepancies between activation of cell membrane stress response and DAP susceptibility have been previously described by our group in *E. faecalis* (30). Indeed, using a clinical strain pair of DAP-S and DAP-R *E. faecalis* recovered from a patient with a fatal bacteremia (16), we showed that introduction of a single amino acid substitution in LiaF (a member of LiaFSR system) resulted in activation of cell membrane remodeling and DAP tolerance in time-kill assays despite minimal changes in the DAP MIC (30). Our results in this work add more evidence to the fact that DAP MIC determination has major limitations to predict actual susceptibility to this antibiotic and support our premise that more accurate tools are critically required.

Using whole genome sequencing, we explored the possibility that variations in LiaX detection with our ELISA test were due to sequence polymorphisms. We did not find any correlation between specific sequence variants and the results of the ELISA or MICs, suggesting that the amino acid substitutions seen in LiaX in our heterogenous population of *Efm* acquired from patients with bacteremia do not affect antibody binding to LiaX, although we cannot rule out differences in *liaX* expression between the isolates that may affect our ability to detect LiaX. These initial results are encouraging and suggest that studies with more specific antibodies (e.g, anti-LiaX monoclonal antibody) could further improve the sensitivity and specificity of the LiaX ELISA, eliminating concerns of potential alterations of LiaX structure due to substitutions.

In summary, LiaX in *Efm* seems to be essential for cell viability upon activation of cell envelope stress mediated by members of the LiaFSR system. Although the exact function of LiaX in *Efm* remains to be fully characterized, its involvement in cell membrane and PBP homeostasis suggest that this protein plays a critical role in defending *Efm* from cell envelope acting antimicrobials. Using an ELISA method, we could show that identifying LiaX in invasive clinical isolates of *Efm* has potential to characterize strains in which the cell envelope stress response is activated, as better surrogate marker for DAP susceptibility. Such a test may be of value to overcome the major limitations of DAP MIC determination using current approaches.

## MATERIALS AND METHODS

### Bacterial strains and growth conditions

We used previously characterized reference *Efm* strains (20) as well as 86 *Efm* isolates (21). The reference and clinical strains used in this study are listed in Table S1. All clinical *Efm* isolates were recovered from the bloodstream of patients enrolled in the Vancomycin-Resistant Enterococcal Bloodstream Infection Outcomes Study (VENOUS), a prospective cohort study of U.S. adults with enterococcal bloodstream infection (21). All reference and clinical strains have undergone whole-genome sequencing as described previously (21) prior to the onset of this study. Enterococcal strains were grown on Brain Heart Infusion (BHI) agar or in BHI broth at 37°C with gentle agitation.

### Transcriptional analysis of LiaR

The LiaR regulon of *Efm* R497 and its *liaR* null mutant derivative (R497*ΔliaR*) have been previously described (11, 12) using RNA-seq (31). Quantitative real-time polymerase chain reaction (qRT-PCR) was performed to confirm RNASeq results in relation to the level of expression of the *liaXYZ* cluster. Strains were grown to early exponential growth phase in BHI and RNA extraction was performed using Purelink RNA Mini Kit (Ambion). Three biological replicates from each strain were performed. The samples were treated with Turbo DNAse kit (Ambion) to remove genomic DNA. The cDNA was obtained from ∼1000 ng of purified RNA using SuperScriptTM II Reverse Transcriptase (Invitrogen). Evaluation of gene expression was conducted with 10 ng of cDNA using SsoAdvanced Universal SYBR Green Supermix (Bio-Rad) in CFX96 Touch TM Real-Time PCR Detection System (Bio-Rad). Relative expression ratios were calculated by normalizing to the housekeeping genes *gyrB*. The primer efficiency was calculated using a corrected calculation model described by Pfaffl (32). Primer efficiency was determined by LinRegPCR program for each reaction. Differences in gene expression between the pair of strains were calculated using the normalized expression for each gene. Primers used for qRT-PCR are listed in Table S4.

### Bacterial mutagenesis and complementation

We generated non-polar deletions of *liaR* and *liaX* in strains of *Efm.* Deletion mutants of *Efm* TX1330RF were created using the p-chloro-phenylalanine (p-Chl-Phe) sensitivity counterselection system (PheS*) using pHOU1 plasmid, which confers resistance to gentamicin, as described previously (23). Briefly, upstream and downstream fragments of *liaR* and *liaX* were amplified by PCR using primers listed in Table S4. PCR products were cloned into pHOU1 using BamHI and EcoRI restriction sites. The plasmid constructs were electroporated into *E. faecalis* CK111 and subsequently delivered to *Efm* TX1330RF by conjugation. First-recombination integrants were selected on plates containing gentamicin 150 μg/ml and fusidic acid 20 μg/ml. Subsequently, these integrants were grown on p-Chl-Phe and selected based on gentamicin sensitivity. Gentamicin-susceptible colonies were screened for deletion of the desired gene by sequencing of the entire *liaR* or *liaX* open reading frames. Mutants were characterized by pulsed-field gel electrophoresis (PFGE) and antibiotic susceptibility testing. For transcomplementation, the *liaR* gene was amplified by PCR (using primers in Table S4) using total DNA from *Efm* TX1330RF as template. The resulting fragment was cloned into pMSP3535 (33) under the control of the P*nis*A promoter which allows inducible expression in the presence of nisin. Recombinant pMSP3535 derivatives were purified on Luria-Bertani agar containing erythromycin (100 μg/ml). All inserts were confirmed by Sanger sequencing before introducing into *Efm* TX1330RF derivatives by electroporation. All *Efm* derivatives with pMSP3535 were maintained with erythromycin (20 μg/ml) and expression was achieved with nisin (50 ng/ml). Strains and plasmids used in mutagenesis are listed in Table S5.

### Antimicrobial susceptibility testing

Minimum inhibitory concentrations (MICs) for DAP for all *Efm* strains and isolates were performed via broth microdilution (BMD) and gradient strip. For BMDs, serial two-fold dilutions of DAP were prepared in a 96-well microtiter plate in Mueller-Hinton broth (BD™) supplemented with Ca_2_Cl 50 mg/L. A 0.5 McFarland turbidity standard was prepared per strain, diluted to 5 x 10^5^ CFU/mL, and inoculated into microtiter wells. Broth microdilution assay preparations were conducted by two different researchers. Plates were read by three independent observers after incubation at 37°C for 24 hours. For gradient strip testing, a 0.5 McFarland turbidity standard was prepared per strain and inoculated on Mueller-Hinton agar. After bacterial solution absorption, DAP gradient Etest strips (bioMérieux, Inc., Durham, NC) were applied to the agar surface. After incubation at 37°C for 24 hours, MICs were read at the location where the elliptical zone of inhibition intersected the strip.

### *Expression and purification of LiaX from* Enterococcus faecium

The gene encoding LiaX from R494 was cloned into a modified pET vector and transformed into *E. coli* BL21(DE3). LiaX was overexpressed by growing cells in Luria Bertani (LB) broth containing 50 μg/mL kanamycin and inducing with 0.5mM isopropyl β-D-1-thiogalactopyranoside (IPTG) for 20 h at 16°C. Cells were pelleted and resuspended in buffer A (20mM HEPES pH 7.4, 300 mM NaCl, 20 mM imidazole) with EDTA-free complete protease inhibitor tablets (Roche Diagnostics Corp, Indianapolis, IN) before undergoing lysis by sonication. Proteins were purified using a HisTrap FF crude column (GE Healthcare Life Sciences, Marlborough, MA). The protein was eluted with a continuous elution gradient of 20 to 500 mM imidazole. The LiaX fractions were pooled and the His tag was subsequently removed by TEV protease (34). The flowthrough was then dialyzed against 20 mM Tris-HCl pH 8.0 and purified over a Q-XL Sepharose column using a 0.1 mM to 1000 mM NaCl gradient. The protein was further purified by size-exclusion chromatography, and the fractions containing LiaX were pooled and concentrated. Purification fractions and final pooled purified LiaX was run on 15% sodium dodecyl sulfate polyacrylamide gel electrophoresis (SDS-PAGE), with purity assessed by ImageJ (National Institutes of Health) software.

### Production and purification of polyclonal antibodies to LiaX

The purified LiaX protein was used to raise antisera in goats at Bethyl Laboratories (Montgomery, TX). The antisera subsequently underwent high-affinity antigen-based purification using CnBr-activated Sepharose coupled with purified LiaX protein (GeneMed Synthesis, Inc., San Antonio, TX). Antibodies were eluted with 0.1M Glycine-HCl pH 2.5 and immediately neutralized after elution with 1M Tris, pH 8.0.

### ELISA antibody titer of goat polyclonal antibodies

To assess analytical sensitivity of the purified polyclonal goat antibodies to LiaX, an ELISA titer was conducted. Comparator goat pre-immune sera was used as a control to ensure absence of LiaX-binding antibodies prior to immunization with LiaX, and to monitor for generation of nonspecific background signal due to antibody complexes in the absence of antigen. The ELISA was performed in triplicate in a 96-well microtiter plate with 10 ng purified LiaX per well and donkey anti-goat alkaline phosphatase-conjugated secondary antibody and chromogenic p-nitrophenyl phosphate (PNPP) substrate.

### Western blotting for protein localization and analytical specificity

Strains were grown to mid-exponential phase. For localization assessment, cells were pelleted, and supernatant was aspirated. Supernatants were filtered through a 0.22-μM filter (Millipore, Sigma) then 10% TCA was added and incubated at 4°C overnight. Protein was pelleted from the TCA supernatants at 15,000 rpm for 20 minutes at 4°C. Supernatant protein pellets were washed with 500 µL cold acetone twice, dried under vacuum for 60 seconds, and resuspended in 100 µL of phosphate buffered saline (PBS) with phenylmethylsulfonyl fluoride (PMSF) protease inhibitor before mechanical lysis with glass beads at 4°C using a FastPrep 24 instrument. Cell debris and beads were spun down at 13,000 rpm for 20 minutes at 4°C. The cell lysates were collected, avoiding debris. Supernatant protein and cell lysates underwent bicinchoninic acid protein concentration assay in three biological replicates at 1:10 and 1:25 dilution for preparation of loading 10 μg of protein per SDS-PAGE well. Western blotting was performed on 8% SDS-PAGE gels and transferred to polyvinyldifluoride (PVDF) membranes using the iBlot 2 (Thermo Fisher Scientific). Membranes were blocked and probed with purified polyclonal goat antibodies to LiaX diluted 1:3000 and horseradish peroxidase-conjugated mouse anti-goat antibody diluted 1:3000. Membranes were incubated with SuperSignal West Pico Chemiluminescent substrate (Thermo Scientific) and visualized using a ChemiDoc Imaging system (BioRad). Relative intensities for the 58.6 kDa molecular weight band representing Efm LiaX were determined using the Gel Analyzer tool in Fiji (35) and normalized to the positive control purified Efm LiaX protein band.

### The LiaX ELISA optimization and final protocol

The optimal antibody concentrations were determined using checkerboard titration. About 10^7^ CFUs of several *Efm* isolates with a wide range of LiaX production were coated to 96-well microtiter plate wells. Controls included 10 ng pure LiaX protein and wells without coated cells or other antigens. The dilutions of purified polyclonal goat antibodies to LiaX ranged from 1:100 to 1:4000. The dilutions of donkey anti-goat alkaline phosphatase-conjugated secondary antibody ranged from 1:1000 to 1:6000. The optimal antibody dilutions were determined as the values that yielded the maximum *A*_405nm_ values within the accurate detection range of the spectrophotometer without generation of non-specific background signal in negative control wells. The final LiaX ELISA was performed as follows: *Efm* isolates were grown to early mid-exponential growth phase and normalized to an *A*_600nm_ of 1.0 before cell pelleting and washing with PBS pH 7.4. The cells were then resuspended in 50 mM carbonate-bicarbonate buffer pH 9.6, and 100 μL (∼10^7^ CFUs) were coated to wells in triplicate in 96-well high-binding microtiter plates for 16 hours at 4°C. Coated wells were washed three times with PBS, then blocked with 2% BSA 0.1% Tween20 in PBS for 1 hour at room temperature. LiaX was detected with anti-LiaX purified polyclonal goat antibodies diluted 1:400 in 1% BSA, 0.05% Tween20 in PBS and incubated for 2 hours at room temperature. After 3 washes with PBS, donkey anti-goat alkaline phosphatase secondary antibody was diluted 1:2000 1% BSA, 0.05% Tween20 in PBS, added to the wells, and incubated for 1 hour at room temperature. After 3 final washes with PBS, colorimetric substrate PNPP was added to the wells and incubated in the dark for 15 minutes at room temperature, and then A_405_ nm was read using a plate spectrophotometer. Isolates were tested in at least three separate assays with three technical replicates per assay.

### Analysis of association of daptomycin MICs and the LiaX ELISA

*A*_405_ nm values from the LiaX ELISA were plotted with DAP-R and DAP-S strains in a Receiver Operating Characteristic (ROC) curve. The cutoff was determined by maximizing the Youden Index. Isolates were plotted according to MIC and positive versus negative LiaX ELISA.

### LiaX and LiaR sequences and association with daptomycin MICs and the LiaX ELISA

The amino acid sequences of LiaX and LiaR (21) for each isolate were acquired. Amino acid sequences were aligned to the consensus sequence of *Efm DO* using the alignment algorithm MUSCLE to determine amino acid substitutions (36). Amino acid sequences for LiaR and LiaX were then analyzed using the R statistical software with each amino acid position as a unique variable. A recursive partitioning model was used to compare each amino acid substitution as a variable in sequential decision trees to compare DAP MICs separately and *A*_405_ of the LiaX ELISA for each isolate. To assess phylogeny of LiaX, maximum likelihood trees were constructed with LiaX protein alignments using RAxML(37) with the PROTGAMMAUTO model.

## ACKNOWLEDGEMENTS

This study was supported by NIH/NIAID grants K24AI121296, R01AI134637, R01AI148342-01, and P01AI152999 to CAA and R01 AI080714 to YS. DBA was supported by an NIH/NIAID T32 fellowship (T32AI141349), NIH Loan Repayment Program award L30AI154520, and Houston Methodist Clinical Scholars Award. SRS was partially supported by NIH/NIAID pre-doctoral training grant T32AI055449. KSH was supported by an NIH/NIAID T32 fellowship (T32 AI141349). BMH was supported by NIAID K01AI148593-01 and P01AI152999. We thank Mauricio Latorre and Audrey Zhao for technical assistance.

